# Cell-Free Optimized Production of Protoporphyrin IX

**DOI:** 10.1101/2023.12.28.573540

**Authors:** David C. Garcia, John P. Davies, Katherine Rhea, Marilyn Lee, Matthew W. Lux

## Abstract

**Background:** The asymmetric and aromatic structures of porphyrins enable semiconducting properties allowing them to absorb light to initiate complex photocatalytic activity. These properties coupled with biological origins have garnered these molecules and their derivatives wide interest as a biotechnological platform. However, porphyrin production in cells is challenging due to the limited titer, long production timescales, and difficult purification.

**Results:** A cell-free metabolic engineering platform was constructed to produce protoporphyrin IX (PPIX) from *E. coli* crude extracts. Using acoustic liquid-handling, design of experiments for high-throughput buffer optimization and co-culturing techniques for extract production, our cell-free reactions effectively produced 0.109 mg/mL quantities of porphyrins.

**Conclusions:** The use of cell-free metabolic engineering as a bioproduction platform could improve the production of toxic or inefficient biomolecules such as PPIX. The engineering strategies including DOE and co-culturing applied in this study provide a roadmap towards increasing the scale of cell-free metabolic engineering by elucidating batch to batch variability, as well as the need for batch specific optimization of reaction conditions.

## Introduction

Porphyrins are molecules composed of one or more cyclic tetrapyrroles whose aromaticity enable semiconductor-like properties making them useful in a broad range of applications, including artificial photosynthesis and light harvesting, catalysis, single-molecule electronics, sensors, nonlinear optics, and chemical warfare agent degradation ^1–6^. Chemical syntheses and isolation of porphyrins from living cells using organic extraction or enzymatic hydrolysis is complex and poses significant challenges for scaling their bioproduction^7–9^. One potential solution to this limitation is through the use of cell-free metabolic engineering (CFME), where enzymatic pathways are reconstituted outside the cell. CFME offers an excellent opportunity to biologically produce molecules with complex metabolic pathways or difficult to implement growth regimes. Removing biological production from a cellular context offers significant advantages as toxic products, deleterious growth conditions, and even lethal metabolic states can be implemented without the need to maintain cellular viability^10,11^. Moreover, CFME can enable far higher throughput than genetic manipulation of cells. While some CFME work uses purified enzymes, here we focus on overexpressed enzymes in crude lysates due to the partially intact cellular metabolism, simpler workflow, and reduced expense. In addition, applications in healthcare settings incentivize protoporphyrin production outside of a cellular context making cell-free production an even more attractive method for the development of medically relevant molecules^12,13^.

The production of porphyrins can be limited by a variety of factors depending on the context, including complex metabolic regulation, slow enzymatic catalysis, and post-production processing^14–16^. In the case of heme production, the cells require extensive reengineering in order to export the major product^11^. These bottlenecks in their metabolism and isolation showcase an excellent opportunity to produce porphyrins outside of the biological limitations of a cell. However, despite the advantages of CFME and the many bench-scale efforts that have shown its effectiveness as a metabolic engineering tool, the factors affecting the scaling of these systems have yet to be deeply interrogated as high throughput CFME is limited by characterization methods reliant on slow and time-consuming chromatographic methods ^17,18^. Additionally, CFME extract production relies on growing and processing separate extracts for each node in a metabolic pathway, significantly increasing the burden of using extracts for both testing and scaled production. As a result, CFME efforts have, to our knowledge, largely remained at the experimental scale.

In this work we take advantage of porphyrins as both a molecule of interest and as an easily detectable product in a cell-free extract to i) show our ability to produce porphyrins using enriched cell-free extracts, ii) explore consolidating the extract source cells into a single co-culture fermentation in order to limit the need for multiple extract productions, and iii) rapidly generate ideal cofactor and substrate mixtures using DBTL-cycles powered by Design of Experiments (DOE). As a result, we showcase the production of PPIX in cell-free extracts, provide the first analysis of a scaled cell-free biological production regime for a small molecule, and provide insights that will further facilitate an in-depth understanding and use of cell-free biomanufacturing.

## Results and Discussion

The previously-characterized pathway examined here involves seven enzymes for over-production of PPIX: HemA- G **(Fig. 1A)**^6^. Our initial experiments were performed by heterologously overexpressing each of these enzymes in separate *E. coli* cultures and lysing the cells to produce extracts enriched with enzymes for each step in the PPIX pathway, referred to as individually enriched extracts for the rest of the manuscript (**Fig. 1B)**. CFME reactions were assembled by combining individually enriched extracts with substrates and cofactors to reactivate the complete pathway outside of the cell. The reaction mixtures contain ATP, Succinate, Glycine, Pyridoxal 5′-phosphate (P5P), and Coenzyme A (CoA) and were incubated at 37°C. Importantly, we found that the CFME reactions expressing HemA-F and HemA-G both showed measurable amounts of a porphyrin that was identified as PPIX by HPLC with fluorescence detection compared to porphyrin standards However, the concentration considerably decreased in the absence of HEMF indicating its importance towards producing PPIX (**Fig. S1)**. Towards rapid optimization of this pathway, all further experiments were measured using plate reader fluorescence measurements of the entire CFME reaction (EX410nm/EM:633nm) and quantified by standard curve. Though the upstream products of PPIX also fluoresce at these wavelengths, the single PPIX peak in the chromatograms supported the assumption that the fluorescence signal was largely derived from PPIX for the purposes of rapid screening. Individually enriched extracts were mixed in several different combinations and the relative levels of extracts were varied to reveal trends in the production of PPIX fluorescence(**Fig. 1B, Fig. S1)**. None of the individual extracts produced high protoporphyrin levels, yet the removal of certain enzymes from the full set still resulted in some PPIX being produced (**Fig. 1B**). This indicates that some endogenous PPIX pathway enzymes may be active in the *E. coli* lysates, though the amount of PPIX being produced was much less than reactions with the full set of enriched enzymes^19^.

**Fig. 1.**
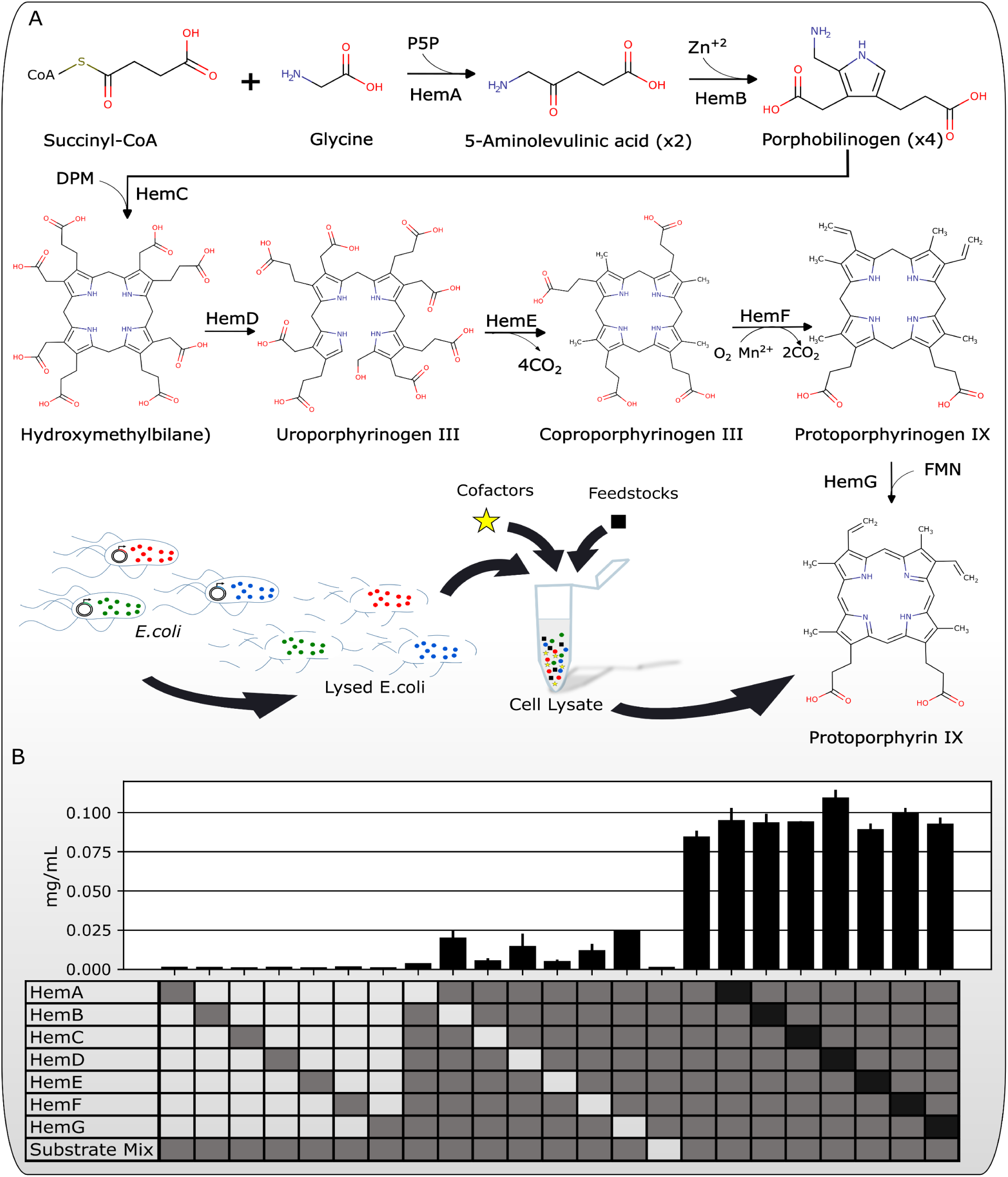
Production of PPIX with CFME. **A.** The C4 pathway for biosynthesis of PPIX. **B.** The full metabolic pathway is reconstituted in cell-free extracts by overexpressing individual enzymes in separate *E. coli* strains and combining enriched extracts in various combinations and relative levels to produce PPIX porphyrins measured using fluorescence (EX410nm/EM:633nm). An empty square indicates the absence of the reagent and black indicates double the concentration. Data for bar plots were acquired using n ≥ 3 biological replicates. Error bars represent standard deviation of the replicates.

While cell-free protein synthesis can be used to produce the enzymes for CFME, the reactions are generally more productive if the enzymes are pre-enriched in cell-extract^20,21^. The labor and cost of producing individual extracts is directly proportional to the number of nodes in the metabolic pathway of interest. In order for CFME to become a viable bioproduction platform at scale for most molecules, the overall cost of the system must be greatly reduced. We decided to explore if it was possible to combine all the steps in the pathway by growing all of the source strains, each expressing only one enzyme, in a co-culture within the same flask, thus allowing for a single fermentation to generate the full metabolic pathway (**Fig. 2A)**. In this scheme, the pathway is optimized with a high degree of control using individual enriched extracts, followed by using a pooled inoculation to scale reactions in a single fermentation.

**Fig. 2.**
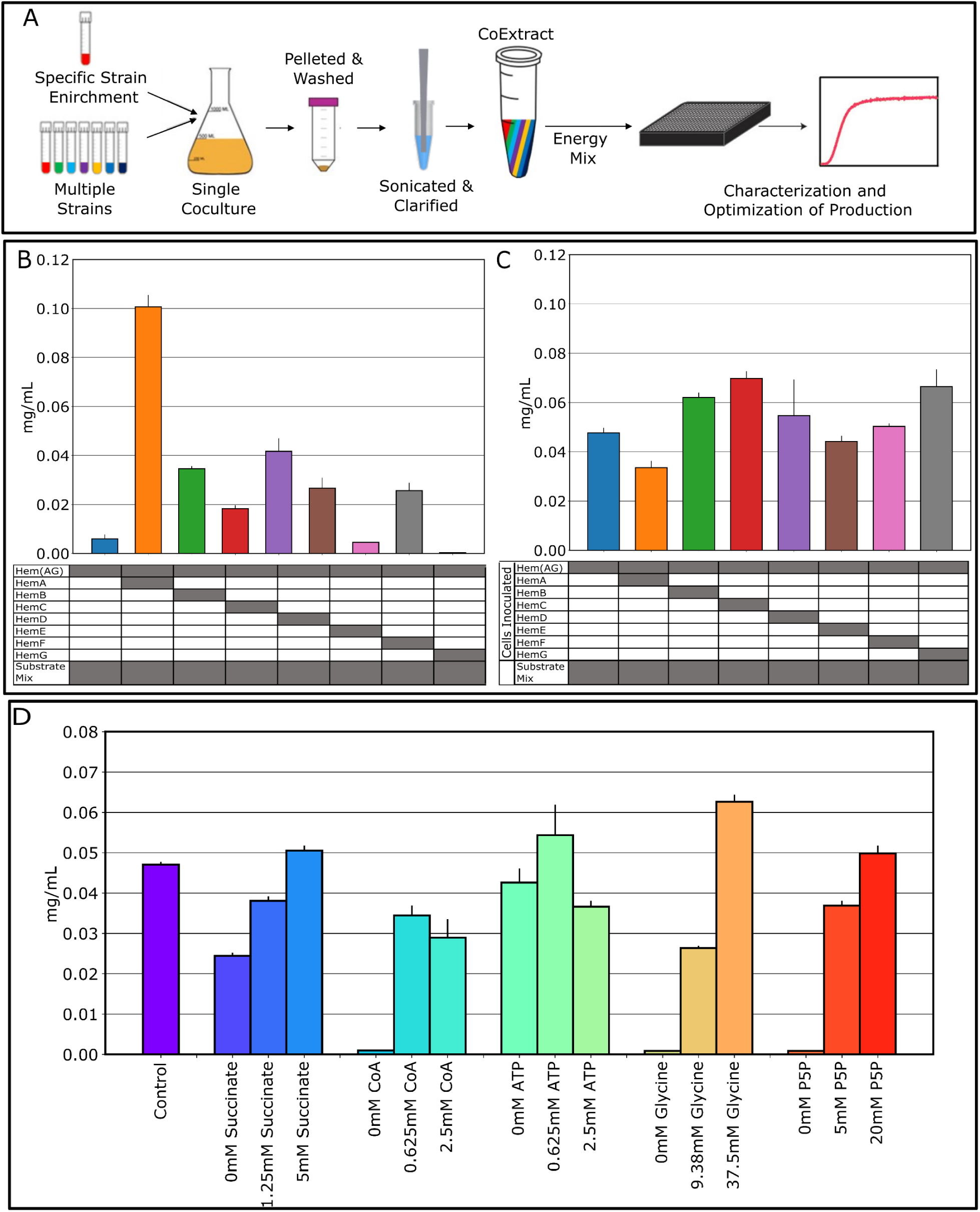
**A.** Cell-free metabolic engineering extract production strategy relying on a single co-culture consisting of all of the strains in the pathway with increased inoculations of lynchpin enzymes. **B.** Productivity of PPIX in CFME reactions created using a base co-culture extract derived from equal inoculation of strains expressing one enzyme HemA-G. The base extract was used at a concentration of 13.5 mg/mL total protein and additional reactions were supplemented with 1 µL individually enriched extract for each pathway enzyme. **C.** Productivity of PPIX in CFME reactions made with co-cultured cell extracts expressing HemA-G, one enzyme expressed per strain, with double inoculation of one strain and equal inoculation of all others. **D.** PPIX production following a cofactor titration in a CFME extract with a double inoculation of HemC. Data for plots were acquired using n ≥ 3 biological replicates. Error bars represent standard deviation of the replicates.

Our first attempt to reduce the number of fermentations required to produce an active CFME pathway consisted of using a base co-culture extract created with equal inoculations of each of the 7 nodes in the PPIX pathway, termed HemAG to indicate the extracts coming from the same culture. This first co-culture lysate had low detected production of PPIX. To troubleshoot, the co-culture HemAG extract was supplemented with an extract of each of the individually expressed nodes (HemAG+HemA, etc.) **(Fig. 2B)**. Supplementing several of the extracts enriched with single enzymes improved the amount of PPIX being produced from the base extract. HemA, the only enzyme not endogenously expressed in *E. coli,* had the greatest effect. These results indicate some strains are being outcompeted in the co-culture, highlighting the limitations of removing the more fine-tuned control of the reaction contents. Replicating the process of producing these extracts as triplicates and with different growth conditions showed the potential for batch-to-batch variability (**Fig. S2**). Though activity in a coculture HemAG extract can be tuned by supplementation with individually enriched extracts, batch-to-batch variation and the natural burden elicited by heterologous protein production is not ideal and will likely require methods of regulating growth to produce a consistent co-culture extract. This is further evidenced by the variability in growth dynamics seen for each of the cells with and without induction. The presence of IPTG significantly impairs the growth of several of the strains, particularly cells carrying the plasmid for HemF, and elicits a general burden on the cells that causes variability within replicates (**Fig. S3**). Though less common in cell-free metabolic engineering applications, batch-to-batch variability in cell-free extracts has been a well-documented factor impacting cell-free protein synthesis^22–24^. Strategies to control the dynamics of cell populations use tools like auxotrophies, or lysis circuits, but impart a further burden on the cells that may limit their bioproduction relevance^25,26^.

We sought to tune growth by increasing the inoculum of strains expressing each node in the pathway. We prepared CFME extracts as before, but with doubled inoculums of each strain containing HemA-G plasmids. We found that the extracts with increased inoculations of HemB, HemC and HemG had improved final protoporphyrin yield (**Fig. 2C)**. However, larger differences between batches are clear when comparing the equal inoculum cases (“HemAG”) across experiments **(Fig. 2B and C, Fig. S2)**. While an increased inoculum of *E. coli* with the HemC plasmid substantially improved the PPIX titer while being one of the slowest growing strains when induced, this trend is not consistent as increasing the inoculum of the HemG strain, one of the fastest growing, reached a similar PPIX concentration. Further work will be required to more effectively measure, predict, and control the exact makeup of the cell-free extract source consortium, as even minor variations in growth could substantially change the proteome of the consolidated extract^27,28^. Despite the need for improved control for reproducibility in co-cultured CFME extracts, promising product yields incentivized us to further explore and optimize these combined reactions.

The HemAG co-culture extract prepared with a double inoculation of HemC, termed HemAG-2xC for the remainder of this work, was used to further explore the effects of optimizing concentrations of CFME ingredients and scaling lysate preparation. The levels of each fed substrate and cofactor can greatly impact productivity, and some of these molecules represent substantial portions of the cost of the CFME reaction (**Table S1**). We performed a set of titrations for each of the components, starting with a HemAG-2xC extract prepared from a 0.2 L shake flask culture volume. The control reaction uses 2.5 mM succinate, 1.25 mM CoA, 1.25 mM ATP, 18.76 mM glycine, and 10 mM P5P. Large changes in PPIX bioproduction occurred when modifying the mixtures and results indicated that for this 0.2 L scale lysate, the CoA, glycine, and P5P were all essential to the function of the system (**Fig. 2D)**. Interestingly, while the expected yield of PPIX from 37.5 mM glycine was only 2.39%, the overall PPIX titer of reactions reached 0.063 mg/mL, comparing favorably with previous efforts producing a similar product^14^. With respect to cost reduction, CoA, ATP, and P5P are the most expensive reagents **(Table S1).** The results showed that ATP could be removed completely to decrease cost without dramatically reducing yields; CoA was required but could possibly be reduced without loss of yield; and P5P concentration increased yield, contrary to cost reduction goals.

Little has been published to date on how CFME reactions perform across scales. To begin to address this knowledge gap, we explored how larger culture volumes would impact the production of CFME lysate and the resulting PPIX product titers. We examined the PPIX productivity of a CFME extract produced from 0.2 and 3 L flask fermentations. In addition to culture volume differences, the 3 L extracts were lysed using a microfluidizer while the 0.2 L culture was processed by sonication. The 0.2 L cultures regularly reach a final OD600 of 5-8, but the 3 L fermentation only reached a final OD600 of ∼3.0. We found that each lysate gave a substantially different PPIX yield using the same energy mixture with higher culture-volume extract reaching only 28% the titer of the 0.2 L fermentation (**Fig. 3A)**.

**Fig. 3.**
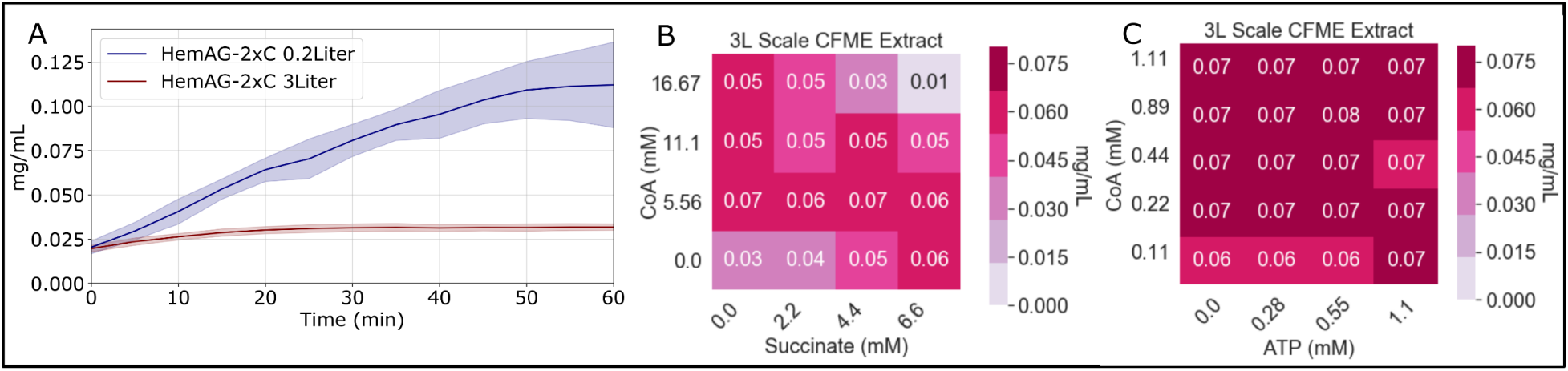
A. Traces of the 3-liter compared to bench scale PPIX reaction. B. Heatmap of CoA titrations with succinate. C. Titrations of reduced concentrations of CoA and ATP.

To further elucidate the differences between lysates, we next evaluated the substrate and cofactor dependence of CFME reactions derived from the 3 L culture. Given that prior results from a 0.2 L extract in Fig. 2D showed limited dependence on ATP and the possibility to reduce CoA concentrations, we titrated these components. We found no clear dependence on either component over the ranges tested for the 3L lysate (**Fig. 3B)**. Interested in this observation, we further titrated the concentration of CoA compared to succinate, finding that neither was essential in the presence of the other for the reaction to function (**Fig. 3C)**. This implied that a cofactor pool was still present. Succinate and CoA could produce the essential precursor, succinyl CoA, through independent pathways, the former by completing a loop through the TCA-cycle and the latter by serving as co-factor in a number of reactions that produce succinyl-CoA. CoA was essential in CFME reactions prepared from 0.2 L scale cultures that reached a higher OD perhaps because cofactors like CoA are depleted at higher OD. Previous work has shown that cofactor pools are significantly different depending on the growth stage of the culture, with cofactor levels in particular depleting heavily over time^29,30^. Though not explicitly measuring cofactors, further evidence that varying growth conditions substantially impacted the metabolism and proteome of a cell-extract was shown with changes to the media and the OD at harvest^31^.

Since there are so many factors to explore in the design of CFME reactions, we decided to apply a Design of Experiments (DOE) approach to more quickly find optima and identify trends the potential search space using a design-build-test-learn (DBTL) cycle (**Fig. 4A)**. Design of Experiments (DoE) is a statistical multifactorial approach to both design and analyze an experimental process. DOE experiments modify a number of factors simultaneously and measure the resultant effect on the system. DOE provides an excellent tool to rapidly define the ideal reaction compositions and develop robust and economical cell-free bioproduction platforms. We aimed to maximize the amount of PPIX produced using the 3L scaled-CFME extract by creating an initial exploratory model modifying the relevant cofactors and substrates, specifically succinate, CoA, ATP, Glycine, and P5P to produce a predictive model of the interactions. An I-optimal design composed of 300 experiments capable of estimating linear blending effects and non-linear blending effects between the substrates and cofactors produced several combinations capable of activating PPIX production (**Fig. 4B**) Our model showed that Glycine and P5P were overwhelmingly the most important reagents with P5P having linear blending effect nearly double that of Glycine **(Table S3)**. Each reagent had a positive effect linearly, but with decreasing value as concentrations increased. More complex multi-reagent synergies were found through the DOE to explain the variability in the model used to produce optimal reactions mixtures **(Table S4)**. Specifically, variations containing only CoA, glycine and P5P were found to have nearly 3- fold greater effect on production compared to linear effects and a 4-fold greater effect compared to similar three component combinations **(Table S5)**. Overall only twelve of the forty sources of variability in the model were found to have statistically insignificant effects, indicating that complex network underlies cell-free metabolism in regards to PPIX production.

**Fig. 4.**
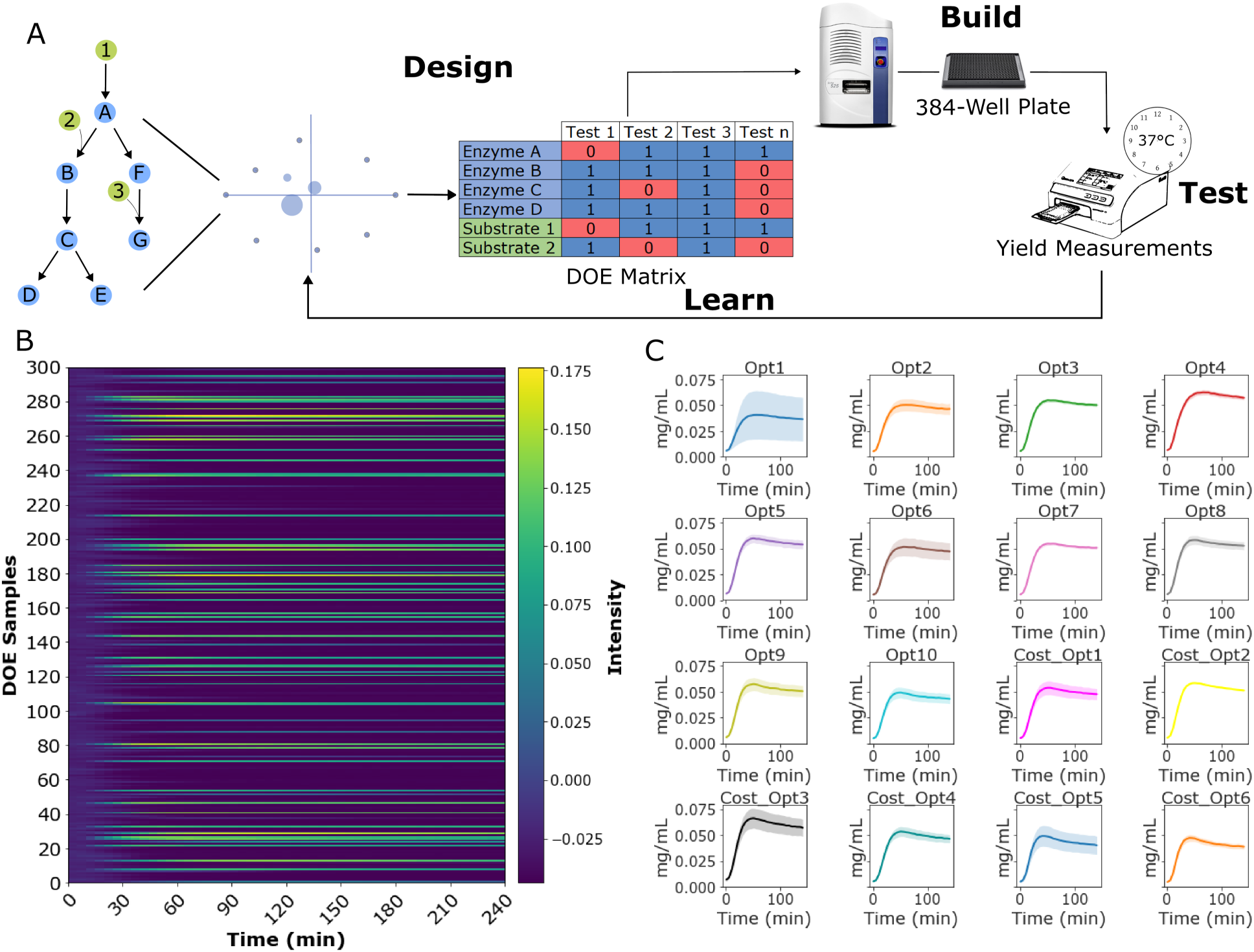
**A.** Graphic illustration of the DBTL cycle used to explore the combinatorial space of CFME compositions. Initial tests of cofactors and substrates defined an initial DOE matrix that was tested, and the resulting data used to define a predictive model for active and optimal reagent concentrations. **B.** 300-experiment DOE heatmap of PPIX produced from the addition of varied cofactors to a HemAG(2xC) CFME extract. **C.** Predicted optimal mixtures for both performance and cost were measured for PPIX production.

Having developed a model that effectively captured the effect of our reagents on PPIX production, we decided to test the DOE model and picked 10 predicted optimal reaction conditions using two objective functions (**Fig. 4C**). The first objective function was set to maximize the production of PPIX (mg/mL) while the second was set to maximize production at the lowest cost($/mg) (**Table S5**). All the predicted optimal reactions yielded final titers greater than 0.03 mg/mL aside from 4 of the cost-optimized reactions that had no PPIX production. Interestingly, in both cases the need for CoA was removed without much change to the overall yield of the reaction. As has previously been noted, changes in growth conditions can have significant impacts on the proteome and resultant metabolome of a cell lysate^11,31^. Additionally, draining cofactor pools could have a substantial effect on the overall function of the extract and indicate why cofactors with large internal pools earlier in the growth, such as CoA would not need to be supplemented^29^. Overall, the highest titer presented in this work of 0.109 mg/mL resulted in a cost of $14.26/mg. The application of the DOE reduced the cost considerably with the best reaction reaching a cost of $1.41/mg, though a lower titer (0.049 mg/mL) was reached (**Table S5**).

### Conclusions

As CFME systems are implemented to produce an expanding range of molecules, an understanding of the underlying mechanisms that control their productivity will need to expand in kind. In this study, the seven-member PPIX synthesis pathway was explored, both to improve the prospects of bioproduction for this interesting molecule, and to uncover factors with the greatest impact on CFME productivity. To start, the ability to produce PPIX in a CFME reaction was confirmed by mixing seven individually enriched extracts for each enzyme in the pathway.

Following successful identification of the product, we pursued two approaches to improve productivity and reduce cost. The first approach examined a co-culture method to produce a multi-strain CFME extract with a single fermentation. We found co-culture cell-extracts both produced PPIX and could be supplemented with bottleneck enzymes post-lysis to improve the overall yield. We further saw that increased inoculums of cells with bottleneck enzymes pre-lysis could effectively increase PPIX titers. Though removing substantial amounts of labor as fewer fermentations are required, very high levels of variability both in the growth of the individual strains and the resultant product titers, incentivize further process improvements to maintain stable communities such as using antibiotics during growth, engineering auxotrophies to limit loss of community members, and chromosomal integration to reduce burden on the cell.

The second approach examined the effects that larger fermentation volume and growth conditions had on the resultant extract. Finding that the cofactor and substrate requirements had changed substantially from a 0.2 to 3-liter scale lysate preparation, we applied a high throughput DOE DBTL cycle to the optimization of the cofactors and substrates required for the reaction. We not only saw that the DOE model could accurately predict an optimal productivity, but it could also select for lower cost reactions using a DBTL turnaround of less than 24-hours. Though the reactions from the 3-liter extract did not reach the same titer as those assembled with individually expressed-enzyme extracts, the final cost of each reaction was substantially lower both in terms of reagents and labor as only a single extract needed to be prepared compared to the 7 required for the highest yield seen in this work. These results shed light on challenges to control both enzyme and small molecule content in different lysate preparations for CFME. At the same time, direct supplementation of additives can quickly be optimized to partially compensate for variability and reach high titers compared to cell-based bioproduction. Importantly, the DOE based optimization of these reactions showed the importance of non-linear effects that are inherent in a complex interconnected system like cell-free metabolism that cannot be explained with simple single reagent titrations. Further analysis of the important synergistic interactions, specifically those with non-linear blending effects while outside the scope of this study, could provide important insight as to the mechanisms that control cell-free metabolism.

The work demonstrated here establishes the foundation for a comprehensive development and understanding of the principles that control scaled production of extracts for cell-free bioproduction and cell-free metabolic engineering. In future work we hope to make use of these high-throughput metabolic perturbations to analyze not just PPIX production, but use it as potential proxy for understanding cell-free metabolism more generally. We expect that expanding on our efforts by directly correlating in-depth metabolic and proteomic analysis to the resultant phenotype of the extract will substantially improve our efforts to use cell-free metabolism as an economical production platform^32,33^. These efforts, together with engineered chassis designed to increase fluxes, improved reaction conditions, and DBTL cycles to optimize reaction formulas; will enable cell-free bioproduction strains to be optimized to meet the technical and economic benchmarks for industrial biomanufacturing.

## Methods

### Cell-Free Extract Preparation

Cell extracts were prepared from *E. coli* BL21(DE3)pLysS cells transformed with one of seven plasmids to express PPIX synthesis pathway enzymes. Plasmids used to produce HemA-F were assembled using a pY71 backbone expression vector containing a T7 promoter, and kanamycin resistance cassette; HemF was synthesized using TWIST Biosciences and expressed using pTwist Amp High Copy. The *hemA* gene from *Rhodobacter capsulatus* (GenBank accession number X53309) was purchased as a gene fragment, then inserted into the PCR-linearized plasmid backbone using NEBuilder HiFi DNA assembly^6^. Sequence information is provided in the supplemental material (**Table S5)**. Cells were grown at 37 °C in 2xYPT (16 g L-1 tryptone, 10 g L-1 yeast extract, 5 g L-1 NaCl, 7 g L-1 KH2PO4, 3 g L-1 K2HPO4). Unless otherwise noted, cell extracts were prepared by seeding with 2.5% v/v of overnight culture and inducing with IPTG at 1 mM at an OD600 of 0.6-0.8. 200-mL or 1 L cultures were grown in 500 mL or 2 L baffled Erlenmeyer flasks, respectively. Cells were harvested by centrifugation at 5000×g for 10 min and washed with S30 buffer (14 mM magnesium acetate, 60 mM potassium acetate, and 10 mM Tris-acetate, pH 8.2) by resuspension and centrifugation. The pellets were weighed, flash-frozen, and stored at −80 °C. Extracts were prepared by thawing and resuspending the cells in 0.8 mL of S30 buffer per gram of cell wet weight. The resuspension was lysed using 530 J per mL of suspension at 50% tip amplitude with ice water cooling. Homogenization was performed as described previously using a Microfluidizer (Microfluidics M-110P) on a cell suspension (prepared the same as the sonication protocol) using one pass followed by centrifugation^34^. Following sonication or homogenization, tubes of cell extract were centrifuged twice at 21,100×g for 10 min at 4 °C, aliquoted, frozen with liquid nitrogen, and stored at −80 °C.

### CFME Reaction Set-up

PPIX production reactions were carried out at 37°C in 4 μL volumes without shaking. Unless otherwise noted, each reaction contained 2.5 mM succinate, 1.25 mM Acetyl-CoA, 1.25 mM ATP, 18.76 mM glycine, and 10 mM Pyridoxal 5′-phosphate. Unless otherwise noted extracts were added to a final protein concentration of 13.5 mg mL−1 as measured by Bradford assay. Reaction components were directly dispensed into a clear-bottom-384 well plate using an Echo 525 Liquid Handler (Beckman Coulter).

### Analysis and Quantification of Porphyrins

PPIX levels were quantified using a standard curve method using synthesized PPIX purchased from Frontier Scientific Inc. (Logan, UT, USA, P562-9), read with a Synergy Neo2 Multi-Mode Microplate Reader (Biotek) set to an excitation and emission of 410 and 633 nm, respectively. Plate reader experiments were performed using 384- well assay plate (Corning, Kennebunk, ME, 04034, USA) covered with a plate sealer (Thermo Scientific, Rochester, NY, 14625, USA) HPLC analysis was performed using Agilent 1290 Infinity II equipped with a diode array detector (DAD) reading at 410 nm and a BDS Hypersil C18 column 150 × 2.1 mM, 2.4 μm particle size (Thermo Scientific, Waltham, MA, USA, 28102–152130). A mobile phase of A: 0.1% formic acid in ultrapure water, and solvent B: 0.1% formic acid in methanol was used at a flow rate of 0.4 mL/min. Injections of 20 µL were separated by a linear gradient transitioning from 100% solvent A to 100% solvent B over 20 min, followed by 100% B solvent for 10 minutes.

### DOE and Statistical Analysis

DOE designs and models were prepared using Stat-Ease Design-Expert 13 and SAS JMP® Pro 15. DOE data analysis was performed using Functional Data Analysis (FDA) applied via the “Functional Data Explorer” platform within SAS JMP® Pro 15 software^35^. A Functional Principal Components (FPC) decomposition was then applied to the response curves. Optimized reaction conditions were found via the Stat-Ease Design-Expert 13 Numerical Optimization feature SAS JMP® Pro 15 software Prediction Profiler Platform. Further data analysis and plots were prepared using custom python scripts.

## Supplementary Information

## Author’s Contribution

DG, ML, and MWL, designed the study; DG and JD carried out the experiments; KR aided with extract production; DG, ML, and MWL wrote the manuscript; ML and MWL supervised the study and secured the funding. All authors read and approved the final manuscript.

## Data Availability

The datasets used and/or analyzed during the current study are available from the corresponding author on reasonable request.

## Declarations

Ethics approval and consent to participate.

Not Applicable.

## Consent for publication

Not Applicable

## Supplemental Figs

**Fig. S1.**
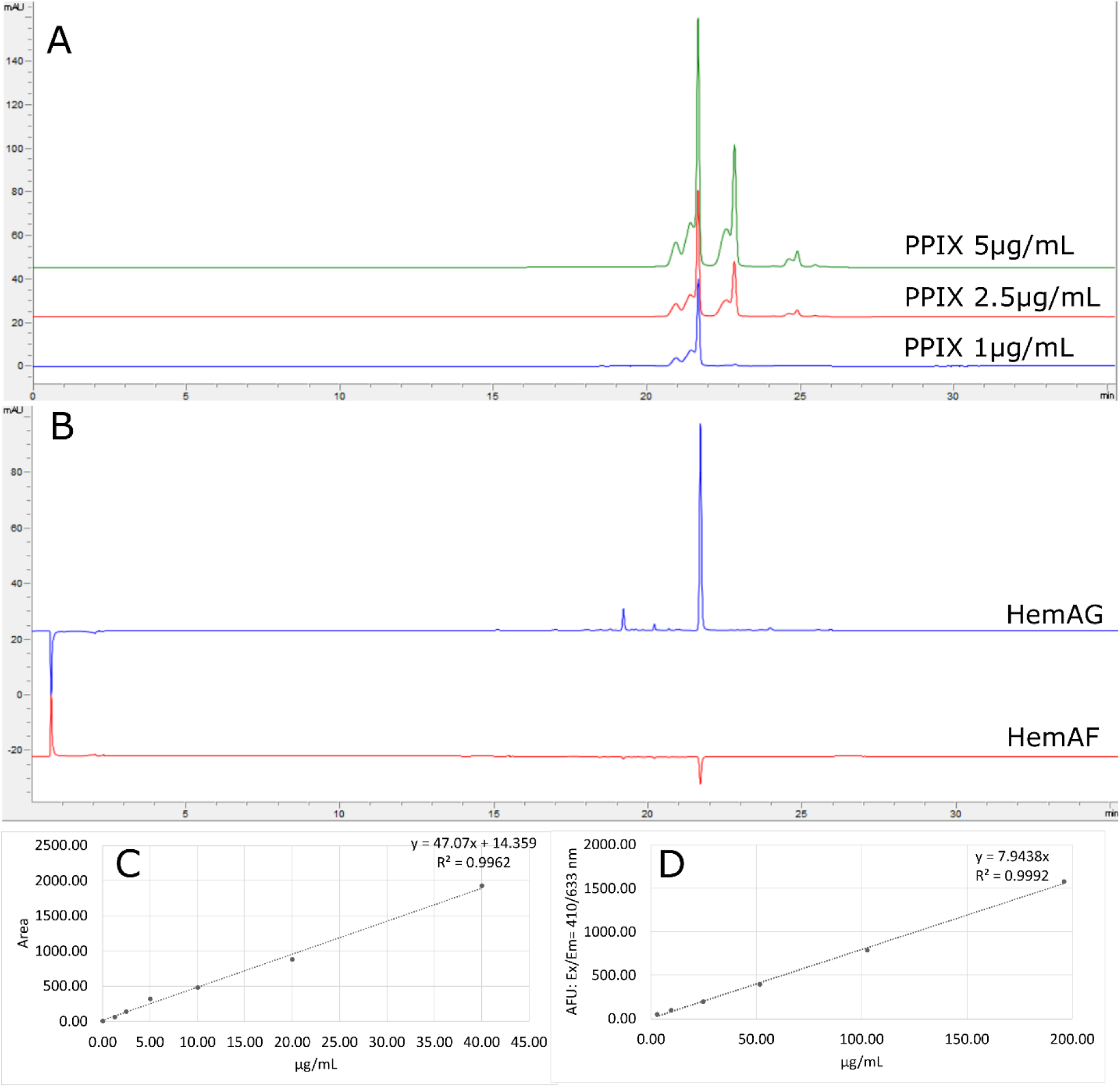
Representative chromatograms of PPIX standards and CFME samples. A. A peak is seen for purified PPIX at 21.6 minutes. B. The same peak is seen at 21.6 minutes when the complete pathway is present in a CFME reaction, but nearly disappears when the last enzyme in the pathway is absent. A small amount of PPIX is detectable as expected without HemG as E. coli naturally produces the enzyme, but not in sufficient quantities to be noticeable by fluorescence measurements. C. PPIX standard curve produced using purified reagent. D. PPIX producing cell-free samples measured with a plate reader, extracted, and cross-referenced to the purified PPIX curve.

**Fig. S2.**
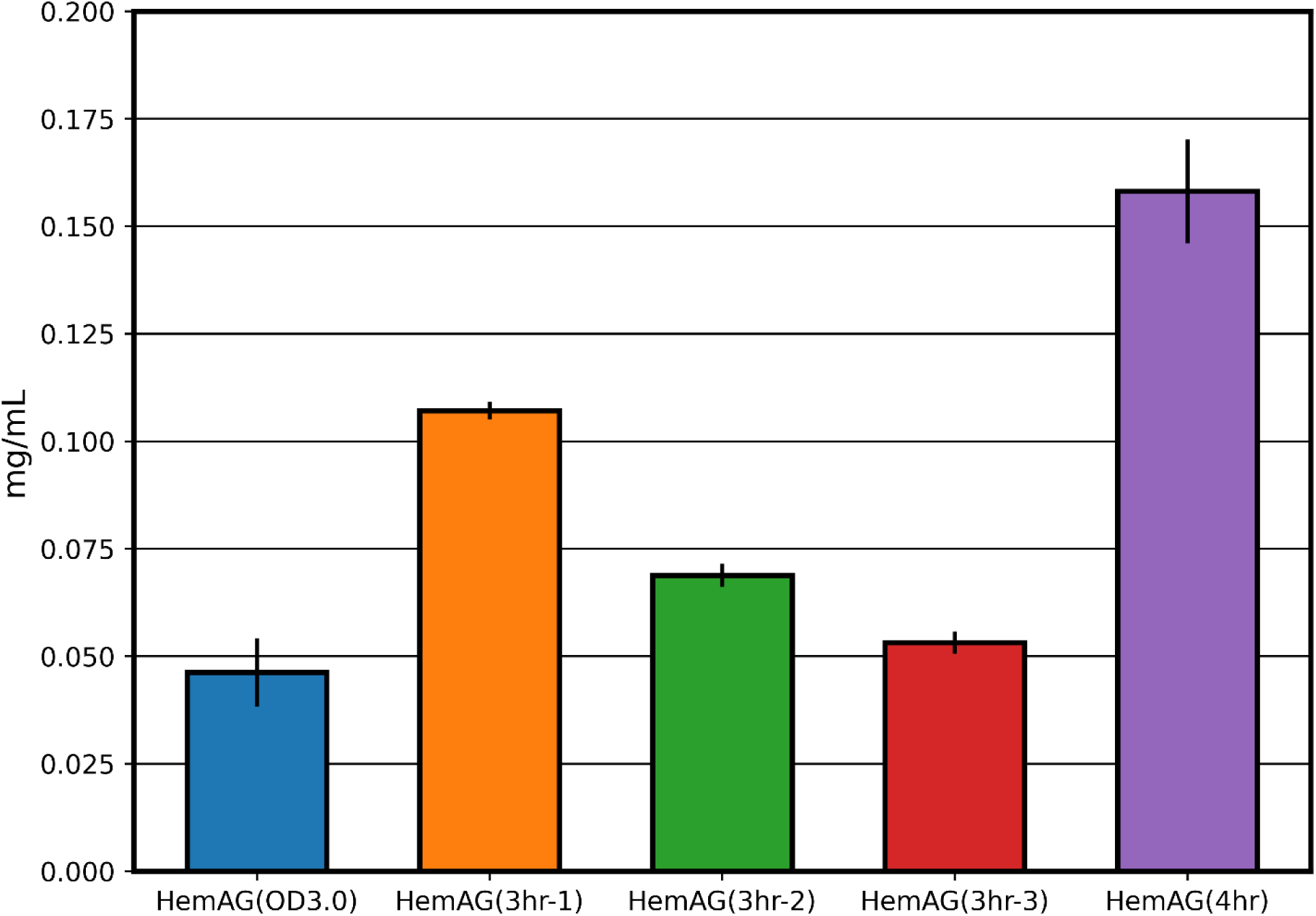
PPIX-producing enriched extracts were prepared using the same inoculation conditions. Following induction with 1mM IPTG, the cells were allowed to grow at 30° C for varying amounts of time or until a specific OD600 as noted in the legend. The CFME reactions were prepared using the conditions noted in the methods.

**Fig. S3.**
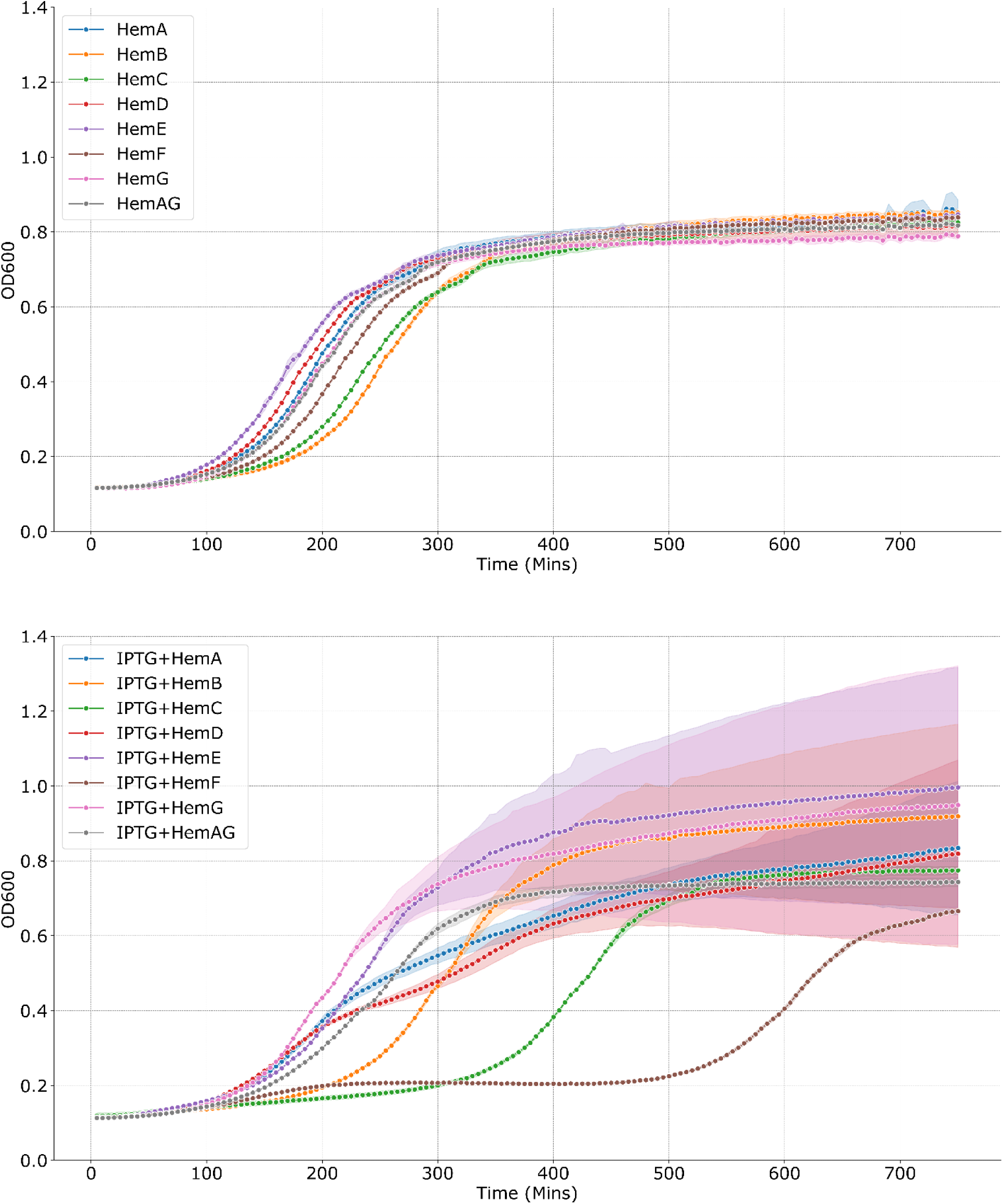
Growth curves of *E. coli* cells expressing PPIX pathway enzymes individually and combined in 2xYPT medium with no antibiotics. **Top:** Cells were inoculated without the presence of IPTG. **Lower**: Cells were inoculated with 1mM IPTG at t=0. Data for line plots were acquired using n ≥ 4 biological replicates. Error bars represent standard deviation of the replicates.

**Table S1:** Reagent costs for cofactors and substrates. Cost based on Sigma Aldrich Checked.

| Reagent | MW | Cost/L(\$) |
| --- | --- | --- |
| Succinate(100 mM) | 118.09 | 0.21 |
| Glycine(500 mM) | 75.07 | 0.16 |
| P5P(100 mM) | 265.16 | 99.91 |
| CoA(20 mM) | 767.53 | 2264.21 |
| ATP(100 mM) | 551.14 | 315.53 |

**Table S2:** Linear blending coefficients acquired from 300-sample exploratory DOE experiment. A positive coefficient indicates the addition of the reagent will improve the production of PPIX.

| Linear blending Coefficient | Reagent |
| --- | --- |
| 12.77 | P5P |
| 6.6 | Glycine |
| 4.49 | ATP |
| 4.39 | Succinate |
| 2.83 | CoA |

**Table S3:**
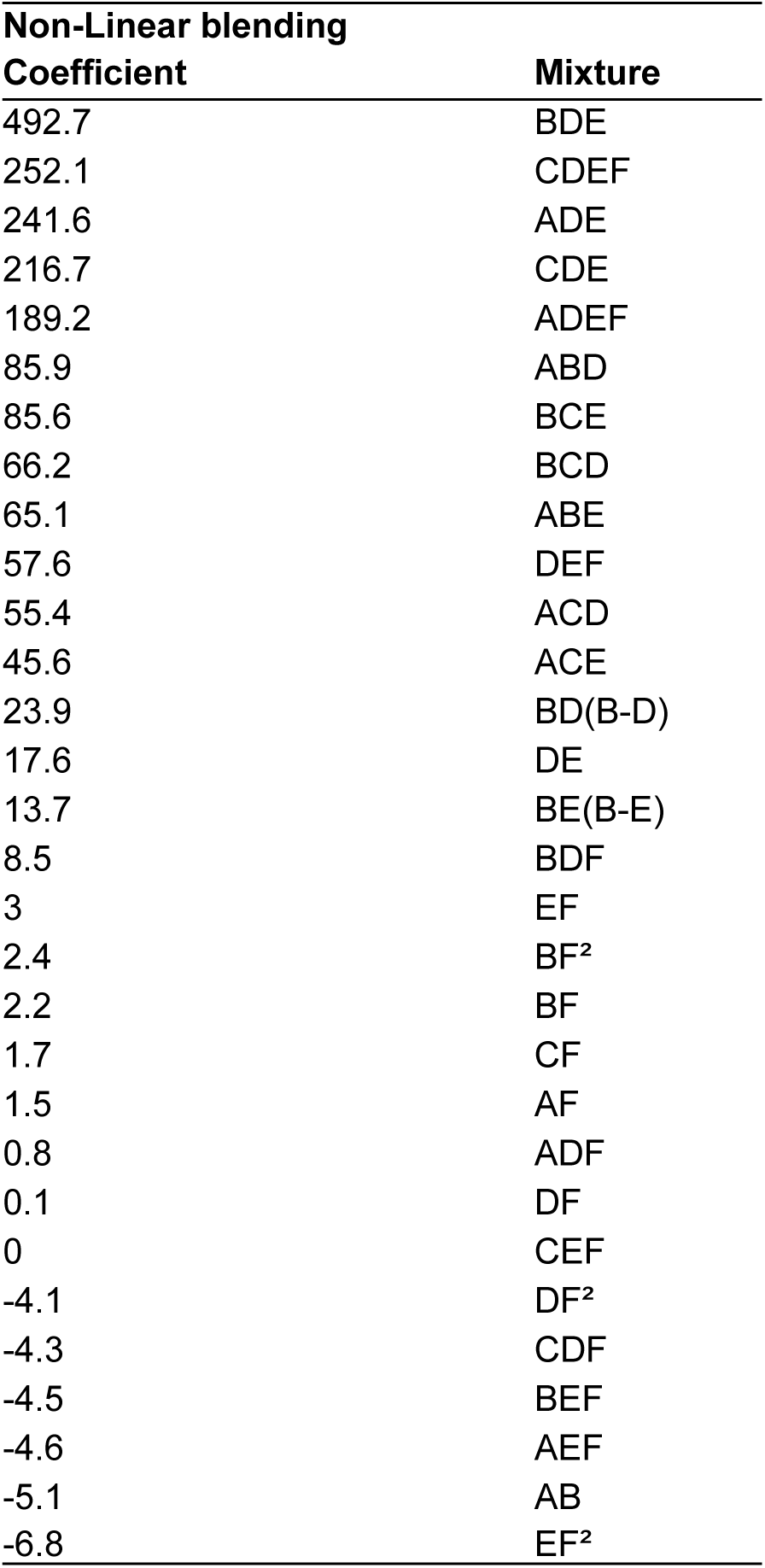
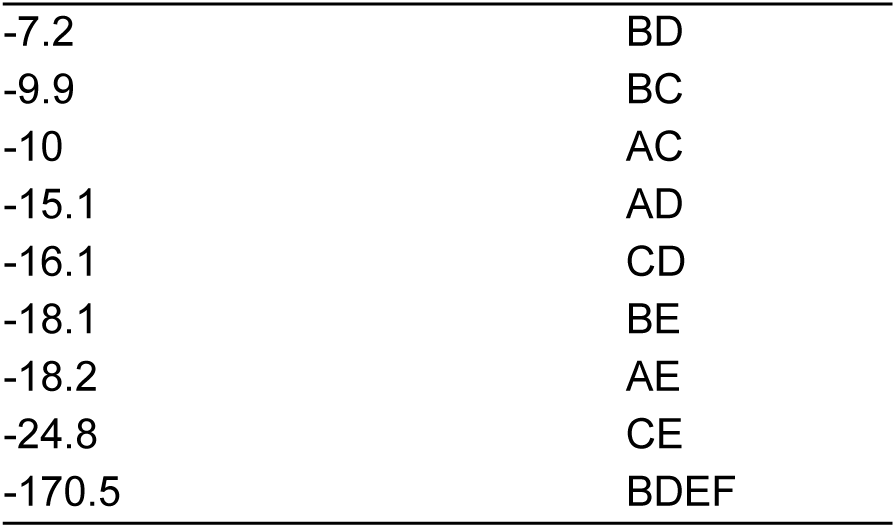
Effects seen though non-linear blending effects of the the different regents and process factors. The letter codes correspond to A: Succinate, B: CoA, C: ATP, D: Glycine, E: P5P, F: total reagent volume.

**Table S4.** Mode terms and effects seen from 44-term model used to produce cost and production optimals using JMP. Source indicates the process factors, interactions, or reagents in the model that contribute to the variability in PPIX production; df indicates the degrees of freedom, and mean square error indicates the variability divided by the number of observations df. The residual value indicates the unexplained variability in the model. The F-value iscalculated by dividing the variability explained by the source by the residual error. The p-values are calculated by the JMP with a value of <0.05 being considered statistically significant

| Source | Sum of Squares | df | Mean Square | F-value | p-value |
| --- | --- | --- | --- | --- | --- |
| Model | 11114.24 | 43 | 258.47 | 64.66 | < 0.0001 |
| Linear Mixture | 1672.76 | 4 | 418.19 | 104.62 | < 0.0001 |
| AB | 5.54 | 1 | 5.54 | 1.39 | 0.24 |
| AC | 22.11 | 1 | 22.11 | 5.53 | 0.02 |
| AD | 38.02 | 1 | 38.02 | 9.51 | 0 |
| AE | 56.83 | 1 | 56.83 | 14.22 | 0 |
| AF | 18.73 | 1 | 18.73 | 4.69 | 0.03 |
| BC | 20.8 | 1 | 20.8 | 5.2 | 0.02 |
| BD | 7.96 | 1 | 7.96 | 1.99 | 0.16 |
| BE | 49.94 | 1 | 49.94 | 12.49 | 0 |
| BF | 38.65 | 1 | 38.65 | 9.67 | 0 |
| CD | 42.98 | 1 | 42.98 | 10.75 | 0 |
| CE | 105.2 | 1 | 105.2 | 26.32 | < 0.0001 |
| CF | 21.64 | 1 | 21.64 | 5.41 | 0.02 |
| DE | 45.59 | 1 | 45.59 | 11.4 | 0 |
| DF | 0.07 | 1 | 0.07 | 0.02 | 0.89 |
| EF | 53.26 | 1 | 53.26 | 13.32 | 0 |
| ABD | 44.63 | 1 | 44.63 | 11.16 | 0 |
| ABE | 24.3 | 1 | 24.3 | 6.08 | 0.01 |
| ACD | 18.33 | 1 | 18.33 | 4.59 | 0.03 |
| ACE | 12.25 | 1 | 12.25 | 3.06 | 0.08 |
| ADE | 298.71 | 1 | 298.71 | 74.73 | < 0.0001 |
| ADF | 0.07 | 1 | 0.07 | 0.01 | 0.89 |
| AEF | 2.22 | 1 | 2.22 | 0.55 | 0.46 |
| BCD | 25.59 | 1 | 25.59 | 6.4 | 0.01 |
| BCE | 41.96 | 1 | 41.96 | 10.5 | 0 |
| BDE | 1233.3 | 1 | 1233.3 | 308.56 | < 0.0001 |
| BDF | 7.12 | 1 | 7.12 | 1.78 | 0.18 |
| BEF | 1.94 | 1 | 1.94 | 0.49 | 0.49 |
| CDE | 242.07 | 1 | 242.07 | 60.56 | < 0.0001 |
| CDF | 1.94 | 1 | 1.94 | 0.49 | 0.49 |
| CEF | 0 | 1 | 0 | 0 | 1 |
| DEF | 222.69 | 1 | 222.69 | 55.71 | < 0.0001 |
| BF^2 | 15.1 | 1 | 15.1 | 3.78 | 0.05 |
| DF^2 | 35.11 | 1 | 35.11 | 8.78 | 0 |
| EF^2 | 99.24 | 1 | 99.24 | 24.83 | < 0.0001 |
| BD(B-D) | 32.52 | 1 | 32.52 | 8.13 | 0 |
| BE(B-E) | 10.68 | 1 | 10.68 | 2.67 | 0.1 |
| ADEF | 75.7 | 1 | 75.7 | 18.94 | < 0.0001 |
| BDEF | 54.21 | 1 | 54.21 | 13.56 | 0 |
| CDEF | 122.66 | 1 | 122.66 | 30.69 | < 0.0001 |
| **Residual** | 923.38 | 231 | 4 |  |  |
| Lack of Fit | 898.59 | 161 | 5.58 | 15.76 | < 0.0001 |
| Pure Error | 24.67 | 70 | 0.35 |  |  |
| Cor Total | 12037.62 | 274 |  |  |  |

**Table S5:** Predicted optimal reaction mixtures from DOE models. Costs were calculated based substrates and cofactors required for each reaction. Reagent concentrations and costs are noted in Table S1.

| **Predicted** |  |  |  |  |  |  |  |
| --- | --- | --- | --- | --- | --- | --- | --- |
| **Optimal Number** | **Succinate(nL)** | **CoA(nL)** | **ATP(nL)** | **Glycine(nL)** | **P5P(nL)** | **Titer(mg/mL)** | **Cost/mg(\$)** |
| Opt 1 | 25 | 0 | 150 | 400 | 425 | 0.04 | 5.91 |
| Opt 2 | 0 | 0 | 175 | 400 | 425 | 0.05 | 5.09 |
| Opt 3 | 50 | 0 | 125 | 400 | 425 | 0.05 | 3.96 |
| Opt 4 | 0 | 0 | 200 | 400 | 400 | 0.06 | 4.31 |
| Opt 5 | 100 | 0 | 75 | 400 | 425 | 0.06 | 2.96 |
| Opt 6 | 25 | 0 | 150 | 425 | 400 | 0.05 | 4.36 |
| Opt 7 | 100 | 0 | 100 | 350 | 450 | 0.05 | 3.68 |
| Opt 8 | 125 | 0 | 50 | 400 | 425 | 0.05 | 2.68 |
| Opt 9 | 175 | 0 | 25 | 375 | 425 | 0.05 | 2.42 |
| Opt 10 | 175 | 0 | 0 | 400 | 425 | 0.05 | 2.39 |
| Cost Opt 1 | 225 | 0 | 0 | 425 | 350 | 0.05 | 1.76 |
| Cost Opt 2 | 200 | 0 | 0 | 450 | 350 | 0.05 | 1.62 |
| Cost Opt 3 | 200 | 0 | 0 | 425 | 375 | 0.06 | 1.56 |
| Cost Opt 4 | 250 | 0 | 0 | 475 | 275 | 0.05 | 1.41 |
| Cost Opt 5 | 0 | 0 | 25 | 550 | 425 | 0.04 | 2.97 |
| Cost Opt 6 | 0 | 0 | 0 | 575 | 425 | 0.04 | 2.62 |
| Cost Opt 7 | 0 | 25 | 0 | 25 | 25 | 0 | N/A |
| Cost Opt 8 | 0 | 0 | 0 | 525 | 0 | 0 | N/A |
| Cost Opt 9 | 1000 | 0 | 0 | 0 | 0 | 0 | N/A |
| Cost Opt 10 | 0 | 25 | 0 | 0 | 0 | 0 | N/A |
| **Base Reaction** | 100 | 250 | 50 | 150 | 400 | 0.11 | 14.26 |

**Table S6:**
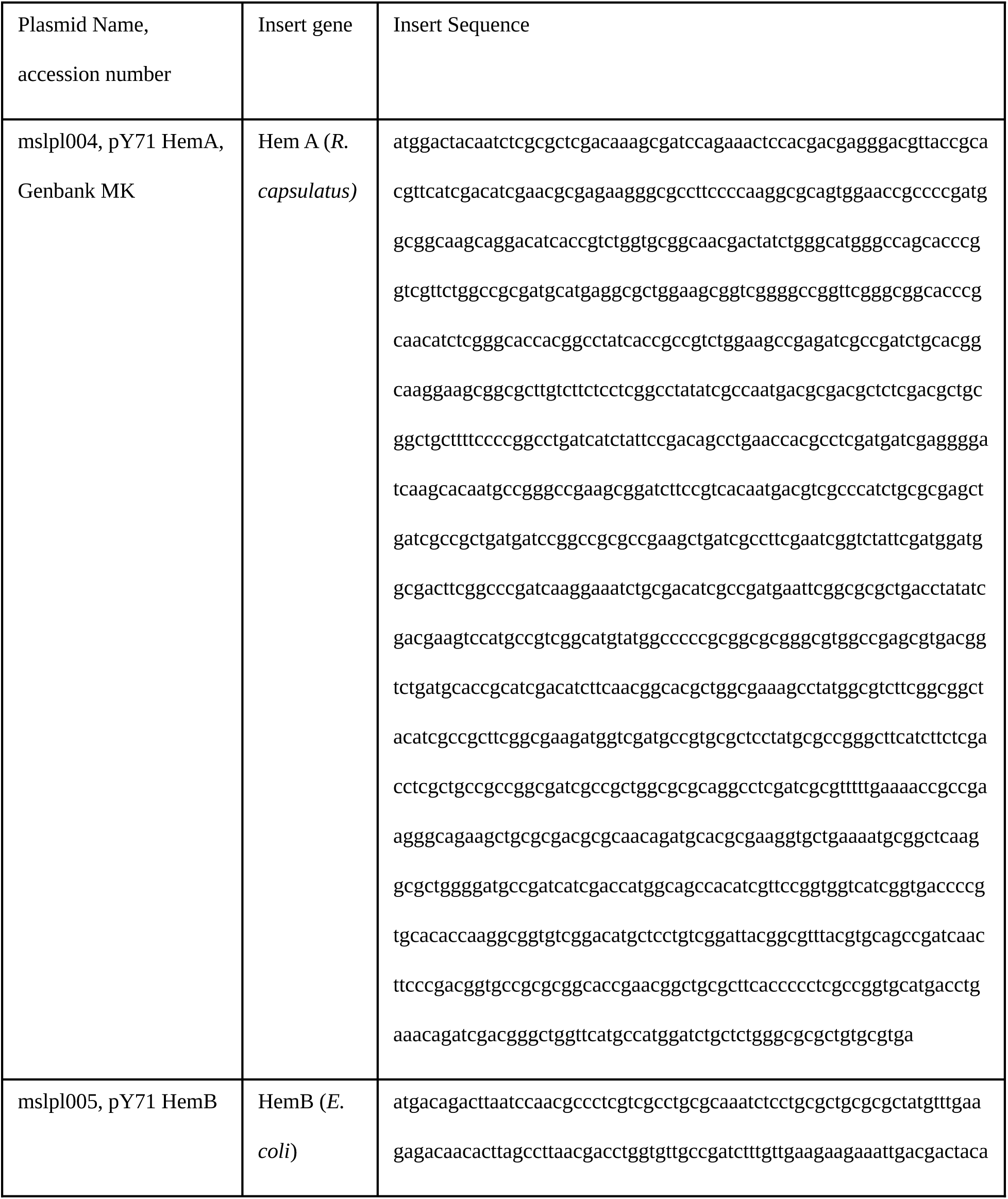
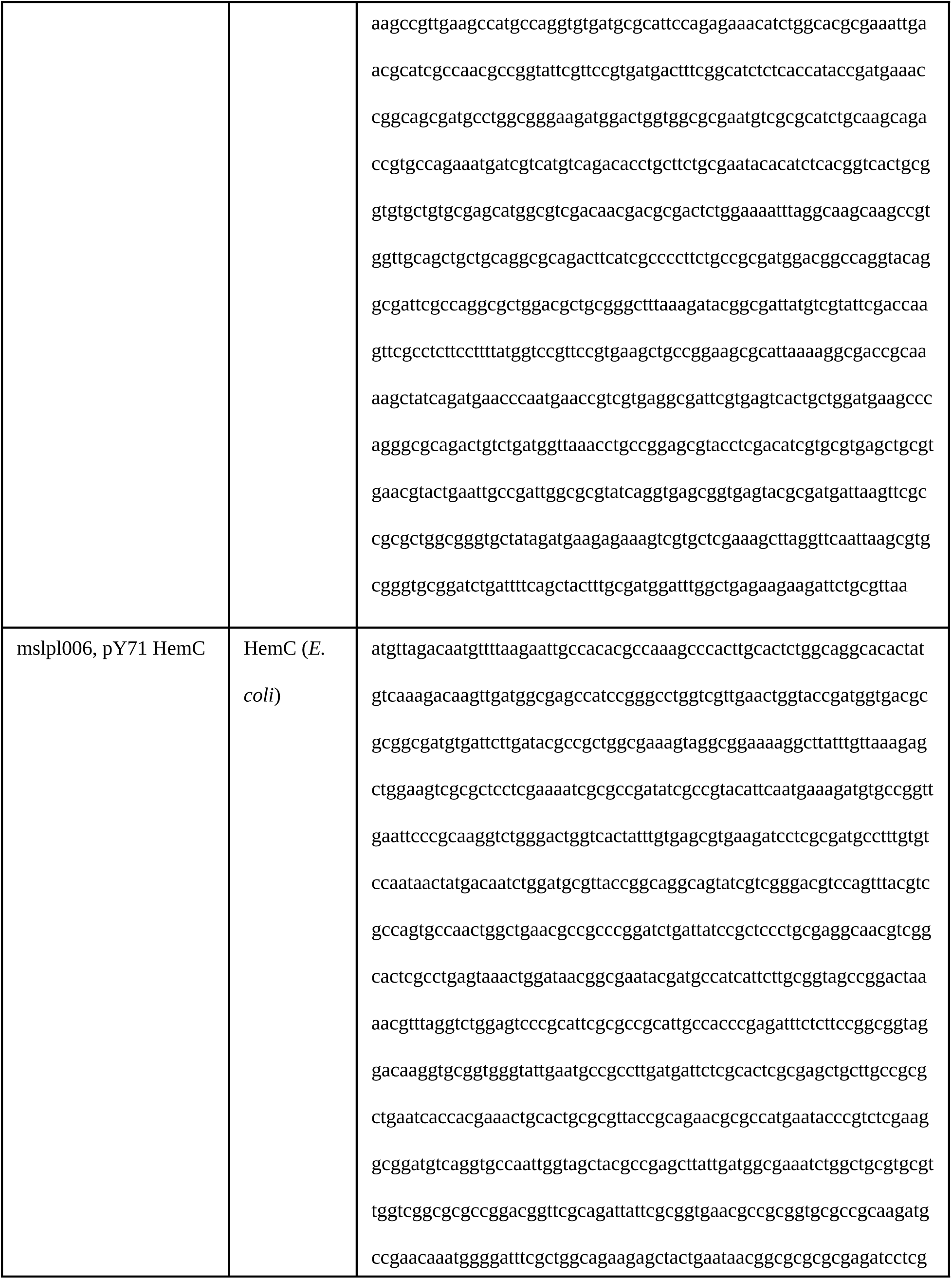
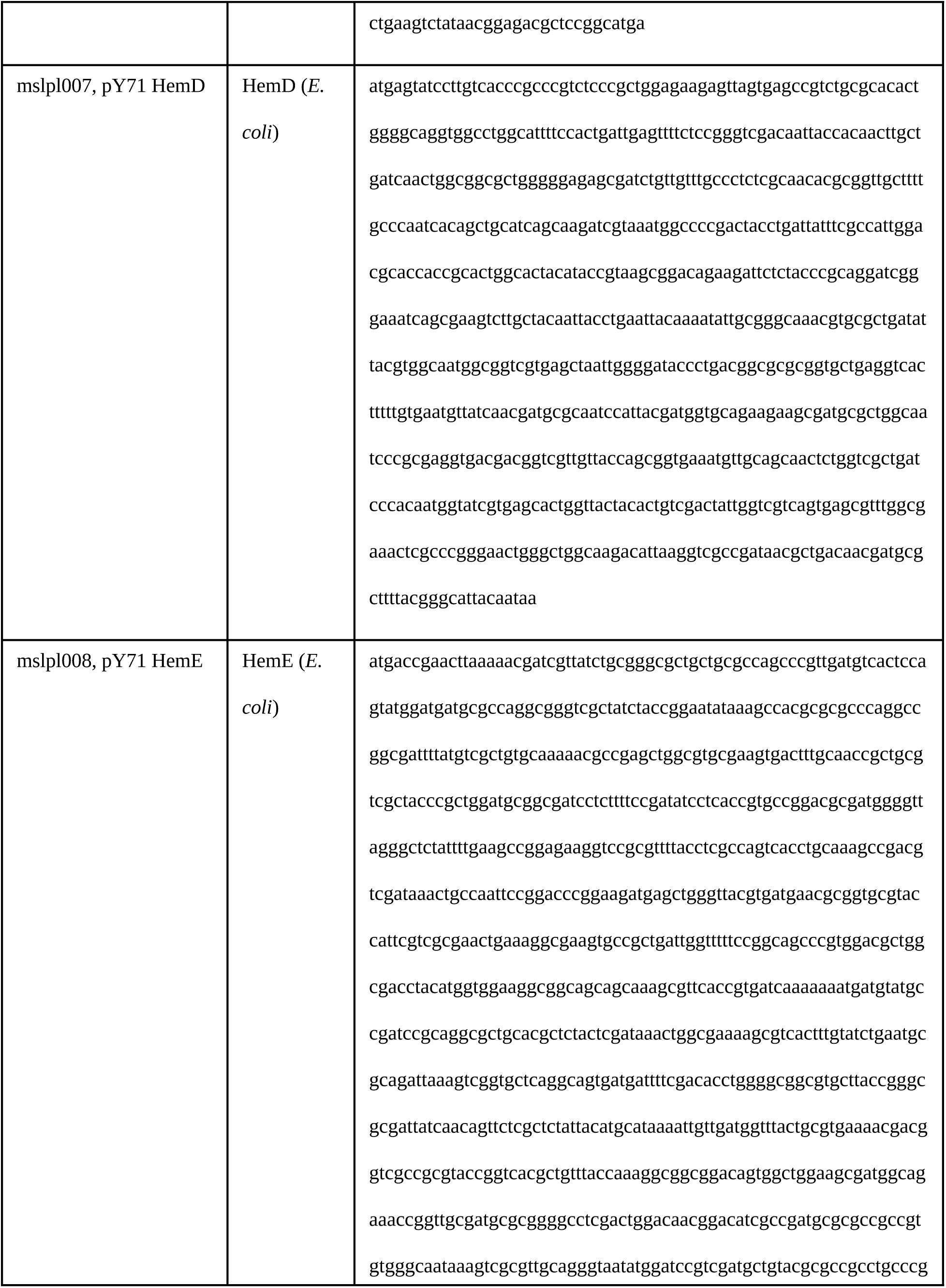
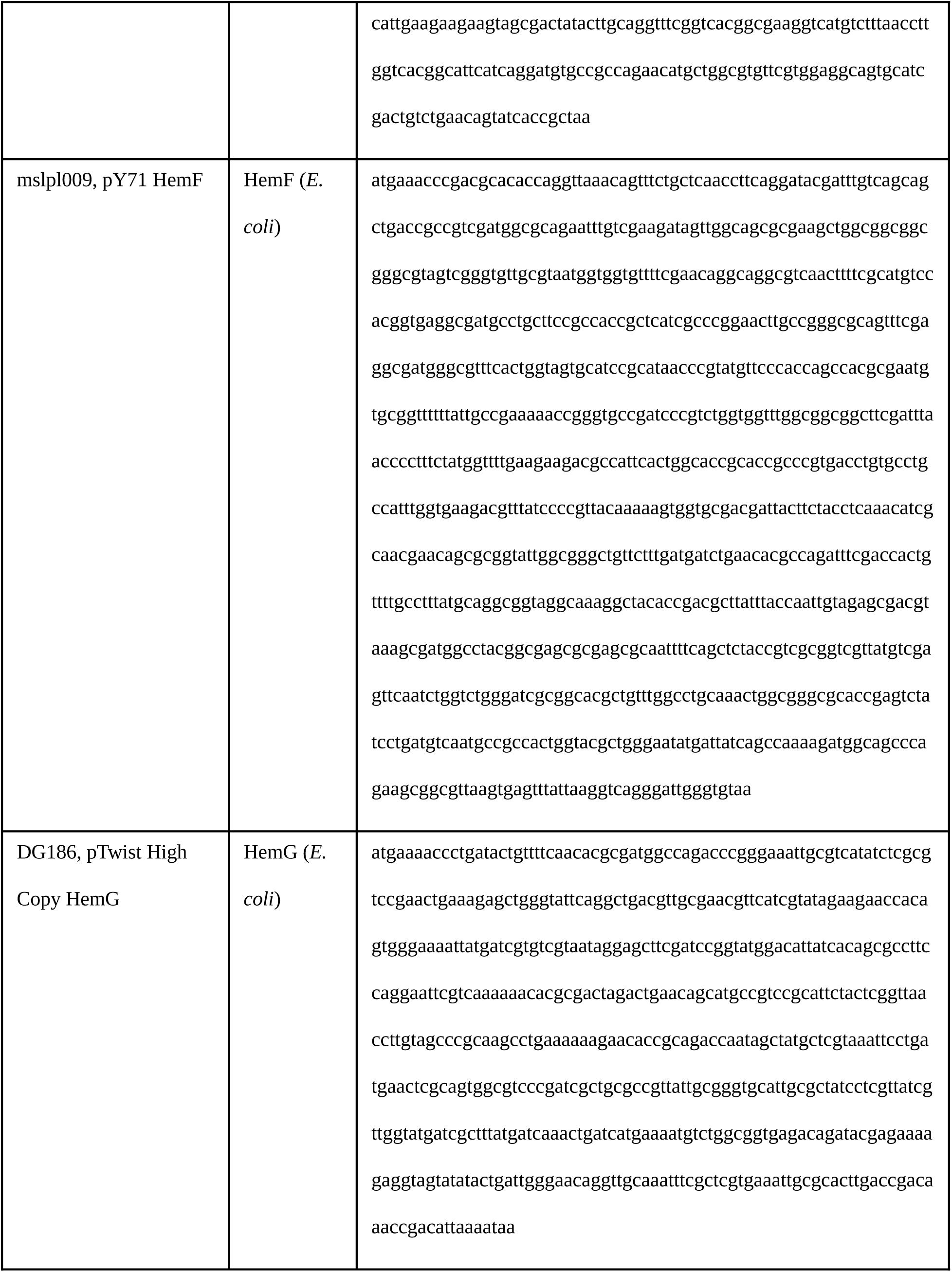
Plasmid sequence information for Hem pathway enzymes.

